# Amotosalen is a bacterial multidrug efflux pump substrate potentially affecting its pathogen inactivation activity

**DOI:** 10.1101/2021.03.15.435562

**Authors:** Alex B. Green, Katelyn E. Zulauf, Katherine A. Truelson, Lucius Chiaraviglio, Meng Cui, Zhemin Zhang, Matthew P. Ware, Willy A. Flegel, Richard L. Haspel, Ed Yu, James E Kirby

## Abstract

Pathogen inactivation is a strategy to improve the safety of transfusion products. The Cerus Intercept technology makes use of a psoralen compound called amotosalen in combination with UVA light to inactivate bacteria, viruses and protozoa. Psoralens have structural similarity to bacterial multidrug-efflux pump substrates. As these efflux pumps are often overexpressed in multidrug-resistant pathogens and with recent reported outbreaks of transfusion-associated sepsis with *Acinetobacter*, we tested whether contemporary drug-resistant pathogens might show resistance to amotosalen and other psoralens based on multidrug efflux mechanisms through microbiological, biophysical and molecular modeling analysis. The main efflux systems in *Enterobacterales* and *Acinetobacter baumannii*, tripartite RND (resistance-nodulation-cell division) systems which span the inner and outer membranes of Gram-negative pathogens and expel antibiotics from the bacterial cytoplasm into the extracellular space, were specifically examined. We found that amotosalen was an efflux substrate for the TolC-dependent RND efflux pumps in *E. coli* and the AdeABC efflux pump from *Acinetobacter baumannii*, and that minimal inhibitory concentrations for contemporary bacterial isolates *in vitro* approached and exceeded the concentration of amotosalen used in the approved platelet and plasma inactivation procedures. These findings suggest that otherwise safe and effective inactivation methods should be further studied to exclude possible gaps in their ability to inactivate contemporary, multidrug-resistant bacterial pathogens.

**Importance:** Pathogen inactivation is a strategy to enhance the safety of transfused blood products. We identify the compound, amotosalen, widely used for pathogen inactivation, as a bacterial multidrug efflux substrate. Specifically, experiments suggest that amotosalen is pumped out of bacteria by the major TolC-dependent RND efflux pumps in *E. coli* and the AdeABC efflux pump in *Acinetobacter baumannii*. Such efflux pumps are often overexpressed in multidrug-resistant pathogens. Importantly, the minimal inhibitory concentrations for contemporary multidrug-resistant *Enterobacterales*, *Acinetobacter baumannii*, *Pseudomonas aeruginosa*, *Burkholderia* spp., *and Stenotrophomonas maltophilia* isolates approached or exceeded the amotosalen concentration used in approved platelet and plasma inactivation procedures, potentially as a result of efflux pump activity. Although there are important differences in methodology between our experiments and blood product pathogen inactivation, these findings suggest that otherwise safe and effective inactivation methods should be further studied to exclude possible gaps in their ability to inactivate contemporary, multidrug-resistant bacterial pathogens.

## Introduction

Bacterial contamination of transfusion product is currently the primary transfusion-related infectious risk (1–3) and is a leading cause of transfusion-related deaths in the United States. Culture-confirmed sepsis is estimated to occur in a least 1 in 100,000 platelet transfusions without pathogen reduction technology (4–6). The need for room temperature storage of platelets contributes to risk by allowing contaminating bacteria to multiply to dangerous levels.

The majority of bacterial platelet contaminants are Gram-positive skin flora. However, recent events highlight potential contamination and transfusion-associated infection with Gram-negative pathogens. In June 2019, the CDC issued a report describing four cases of sepsis attributed apheresis platelet transfusion. Occurring between the months of May and October, 2018, in Utah, California, and Connecticut, the cases were notable for their identification of clonal *Acinetobacter calcoaceticus-baumannii complex* isolates (ACBC). One of these occurred despite use of pathogen reduction technology (7). Following a multi-state investigation, two additional, clonally distinct cases of ACBC platelet-transfusion-associated sepsis were reported in North Carolina, and one additional case was reported in Michigan (7). A summary of blood product contaminants in the United States, the United Kingdom and France published in 2005 found that ∼33% of platelet transfusion-associated infections were caused by either *Enterobacterales* or *Acinetobacter*; the percentage of transfusion-associated infections caused by these pathogens in red blood cell products was even higher at 55% (7, 8). A recent study from the American Red Cross found that *Klebsiella* and *Acinetobacter* spp. were the Gram-negative pathogens most frequently associated with platelet transfusion-associated sepsis (6).

Numerous steps have been taken to protect against bacterial contamination risk. In March 2004, the AABB (previously known as the American Association of Blood Banks) required member institutions to employ means of identifying and mitigating bacterial contamination of platelet products (9). In 2010, the AABB’s standards were updated to recommend pathogen inactivation as well (10). In September 2019, the FDA issued non-binding recommendations to blood collection agencies and transfusion services for pathogen identification, pathogen reduction, and platelet storage (11). In practice, several protocols are now in use for identifying infected blood products including single and pooled unit culture assessment, lipoteichoic acid and lipopolysaccharide antigen detection, and pH-sampling (12–15). Recently, several methods of pre-emptive pathogen reduction, rather than passive detection, have been developed and utilized.

In particular, psoralen compounds have compelling attributes for use in pathogen reduction technologies (16–19) Psoralens are tricyclic, planar compounds capable of forming irreversible, covalent adducts with nucleic acids following excitation with long-wave ultraviolet light (i.e., UVA) (20, 21). Therefore, psoralens can be added to blood products, which are then irradiated to destroy the nucleic acids of any contaminating pathogens.

Psoralen compounds vary in their characteristics. For instance, while the psoralen, 8-methoxypsoralen (8-MOP), has previously been shown to be effective in inactivating many bacterial species, it is less effective in inactivating viral pathogens (22). In comparison, while 4’-aminomethyl-4,5’,8-trimethylpsoralen (AMT) inactivates both bacterial and viral pathogens, it exhibits high mutagenicity in the Ames test, a surrogate for carcinogenic potential, and therefore is considered inappropriate as a blood product treatment (22, 23). An ideal combination of 8-MOP’s safety profile and AMT’s efficacy was found in 4’-(4-amino-2-oxa)butyl-4,5’,8-trimethylpsoralen (trade name, amotosalen) (22–24). In 2014, the Cerus Corporation’s INTERCEPT Blood System, using amotosalen in conjunction with a specialized UVA illuminator, became the first psoralen-based pathogen reduction system licensed by the US Food and Drug Administration for pathogen reduction in platelets (25) and is also now approved for pathogen reduction in plasma (26).

Since amotosalen’s development, multiple studies have reported its efficacy against a broad spectrum of microbial pathogens (27–29). In August 2020, a study supported by the Cerus Corporation also demonstrated inactivation of ACBC and *Staphylococcus saprophyticus* isolated from the transfusion-related sepsis case mentioned above, which occurred despite use of its pathogen-reduction technology. The study showed a >5.9-log reduction in viable pathogen for both isolates after treatment, implying amotosalen susceptibility (30).

Notably, the tricyclic, planar structure of psoralens, including amotosalen, is reminiscent of known bacterial multidrug efflux pump substrates (31). Moreover, increased susceptibility of an *Escherichia coli acrA* mutant to 8-methyoxypsoralen plus ultraviolet radiation was previously noted in 1982, prior to the identification of AcrAB-TolC as the major tripartite efflux pump in *Escherichia coli* (32). With the dramatic emergence of antimicrobial resistance, often associated with overexpression of such efflux pumps, we therefore considered the possibility that multidrug efflux resistance present in Gram-negative pathogens may also confer resistance to amotosalen and related psoralen compounds. To address this possibility, the activity of amotosalen against contemporary, drug-resistant Gram-negative species most commonly associated with blood product contamination, including *Acinetobacter baumannii*, *Escherichia coli*, and *Klebsiella pneumoniae,* were accordingly characterized. The potential of psoralen to act as substrates for major multidrug efflux pumps found in these organisms was also assessed.

## Results

Assessment of amotosalen activity against *E. coli*, *K. pneumoniae*, *A. baumannii*, *Pseudomonas aeruginosa* and *Burkholderia* strains was performed using reference standard minimal inhibitor concentration (MIC) testing. Accordingly, two-fold serial dilutions of amotosalen were prepared in microwell format, and added to bacteria in standard cation-adjusted Mueller-Hinton broth under conditions recommended by the Clinical Laboratory Standards Institute (33). Microplates were exposed to 2 Joules/cm^2^ of UVA, to approximate exposure during platelet pathogen inactivation (17), and incubated overnight to determine the MIC through absorbance measurements as previously described (34–37).

Strains tested were from collections of contemporary multidrug-resistant strains and were generally carbapenem resistant (Table S1). FDA-CDC Antimicrobial Resistance Bank strains are available as a resource for validation of susceptibility testing methods against new and existing antimicrobials (38). Amotosalen MIC varied between and within genera (see Table I, Table S1). Notably, MIC values used in our *in vitro* assay were close to, and for some strains exceeded, the 150 µM concentration of amotosalen used in the Intercept pathogen inactivation procedure in clinical practice. The modal MICs of multidrug-resistant (MDR) *E. coli* and *K. pneumoniae* isolates exceeded the MICs of broadly-susceptible ATCC strains .of the same species; however, broadly-susceptible *A. baumannii* 17978 (39) had an MIC of 128 µM, identical to the modal MIC of MDR *A. baumannii* isolates.

**Table I.** Minimal inhibitory concentration of amotosalen on ATCC quality control strains and multidrug-resistant clinical strains.

| **Species** | **Strains Tested** | **Modal MIC ($\mu\text{M}$)<sup>a</sup>** | **MIC Range ($\mu\text{M}$)** |
| --- | --- | --- | --- |
| *E. coli* 25922 | - | 8 | - |
| *E. coli* strains | 8 | 128 | 64-256 |
| *K. pneumoniae* 13882 | - | 32, 64<sup>b</sup> | - |
| *K. pneumoniae* strains | 8 | 128 | 128-256 |
| *A. baumannii* 17978 | - | 128 | - |
| *A. baumannii* strains | 17 | 128 | 64-256 |

<sup>a</sup>Modal MIC (minimal inhibitory concentration) for individual strains determined from values obtained from 3 separate experiments with two technical replicates per experiment. Modal MIC value for groups of strains were determined from the modal MIC values of the individual strains represented.
<sup>b</sup>Equal number of replicates showed MIC of levels of 32 and 64 $\mu\text{M}$ .

Based on structural similarity of psoralens to known multidrug-efflux pump substrates and frequent elevation of efflux pump expression in MDR Gram-negative pathogens (40), we then examined whether well-characterized, major efflux pumps from *E. coli* and *A. baumannii* were capable of rendering these strains resistant to psoralen compounds. These pumps are classified as RND (resistance-nodulation-cell division) efflux pumps and consist of three components, residing in the inner membrane, periplasm and outer membrane, respectively.

We first used a genetics approach to assess effects on psoralen efflux in isogenic strains that were competent or incompetent for expression of functional *E. coli* RND family efflux pumps. In *E. coli*, the RND efflux pumps depend on the shared outer membrane channel, TolC (41). We therefore compared psoralen MICs in an *E. coli* K-12 parent strain and a TolC-knockout (*ΔtolC*) strain that were otherwise genetically identical (i.e., isogenic strains). For amotosalen, 8-MOP and AMT (see Table II), the MIC in the *ΔtolC* strain was reduced by up to 16-fold, indicating that psoralens are TolC-dependent efflux substrates.

**Tables II.** Effect of a *ΔtolC* deletion, inactivating RND efflux pumps in *E. coli*, on the minimal inhibitor concentration of psoralens.

| **Psoralen<sup>a</sup>** | **Strain<sup>b</sup>** | **MIC (μM)<sup>c</sup>** |
| --- | --- | --- |
| Amotosalen | wt | 32 |
| Amotosalen | *ΔtolC* | 2 |
| 8-MOP | wt | 128 |
| 8-MOP | *ΔtolC* | 32 |
| AMT | wt | 32 |
| AMT | *ΔtolC* | 8 |

<sup>a</sup>8-MOP = 8-methoxypsoralen; AMT = 4'-aminomethyltrioxsalen hydrochloride
<sup>b</sup>*Escherichia coli* Keio wild type (wt) strain and isogenic *ΔtolC* mutant that cannot produce functional RND (resistance-nodulation-division) efflux transporters due to the absence of the required TolC outer membrane component.
<sup>c</sup>Modal MIC (minimal inhibitory concentration) for individual strains determined from values obtained from at least 3 separate experiments with two technical replicates per experiment.

The major MDR efflux pump in *A. baumannii* is the RND, AdeABC pump with each of the three proteins performing analogous roles to those in the tripartite AcrAB-TolC machinery. Multidrug-resistance in *A. baumannii* and other *Acinetobacter* species is commonly associated with upregulation of the AdeABC efflux system (42–44). To examine whether AdeABC could efflux psoralens, we cloned *adeAB* and *adeC* on separate plasmids under control of inducible promoters. These plasmids were then transformed alone or in combination into the *E. coli* AGX100AX strain, which has deletions in the main RND efflux pumps (Δ*acrAB*, Δ*acrEF*) that partner with TolC in *E. coli*, thereby reducing potentially confounding effects from existing efflux pumps in the *E. coli* experimental system (45).

In these strains, we found evidence for pronounced efflux of psoralens, however, only if both the AdeAB and AdeC plasmids were present (see Table III) and induced in the same strain (data not shown), demonstrating the requirement for all three RND components. For amotosalen, the modal MIC of AdeC expressing control strain, with an inoperative pump complex, was found to be 8 µM, which was comparable with the modal MIC of the AG100AX parent *E. coli* strain. In contrast, the AdeABC expressing experimental strain, with an operative, induced pump, exhibited a 32-fold higher modal MIC of 256 µM. This was similar to the high-level resistance of the multidrug-resistant *A. baumannii* AYE strain from which AdeABC was cloned for use in these experiments, which had a bimodal MIC of 128 µM and 256 µM across replicate experiments (Table S1). Although carbapenem-susceptible, AYE is considered a model MDR *A. baumannii* strain and is notable for having caused wide-spread epidemic infection in France with high mortality (46).

**Tables III.** Effect of a heterologously expressed AdeABC efflux pump from *A. Baumannii* on minimal inhibitor concentration of psoralens in *E. coli*.

| **Psoralen Compound** | **Efflux Pump Strain<sup>a</sup>** | **MIC (μM)<sup>b</sup>** |
| --- | --- | --- |
| Amotosalen | pAdeAB + pAdeC | 256 |
| Amotosalen | pAdeC | 8 |
| 8-MOP | pAdeAB + pAdeC | 128 |
| 8-MOP | pAdeC | 16 |
| AMT | pAdeAB + pAdeC | 256 |
| AMT | pAdeC | 8 |

<sup>a</sup>*E. coli* strain AG100AX lacking major endogenous *E. coli* RND efflux pumps ( $\Delta$ *acrAB* $\Delta$ *acrDE*) was transformed with plasmids expressing the indicated *A. baumannii* AdeABC efflux pump components under inducing conditions.
<sup>b</sup>Modal MIC (minimal inhibitory concentration) for individual strains determined from values obtained from at least 3 separate experiments with two technical replicates per experiment.

The AdeB protein is the drug-binding channel and pump, energized by a proton motive force to move substrates form the bacterial cytoplasm into the periplasm (47). We therefore assessed the ability of amotosalen to bind purified AdeB using a fluorescence polarization taking advantage of the inherent fluorescence of amotosalen (see Fig. 1). The dissociation constant (K_D_) for binding to AdeB was 27.9±1.8 µM, in the same range as the 4.9 µM K_D_ of the efflux pump inhibitor, PAβN, for AdeB, and the K_D_ of ethidium bromide (8.7 µM), proflavin (14.5 µM), and ciprofloxacin (74.1 µM) efflux substrates for the homologous, AcrB (31).

**Figure 1.**
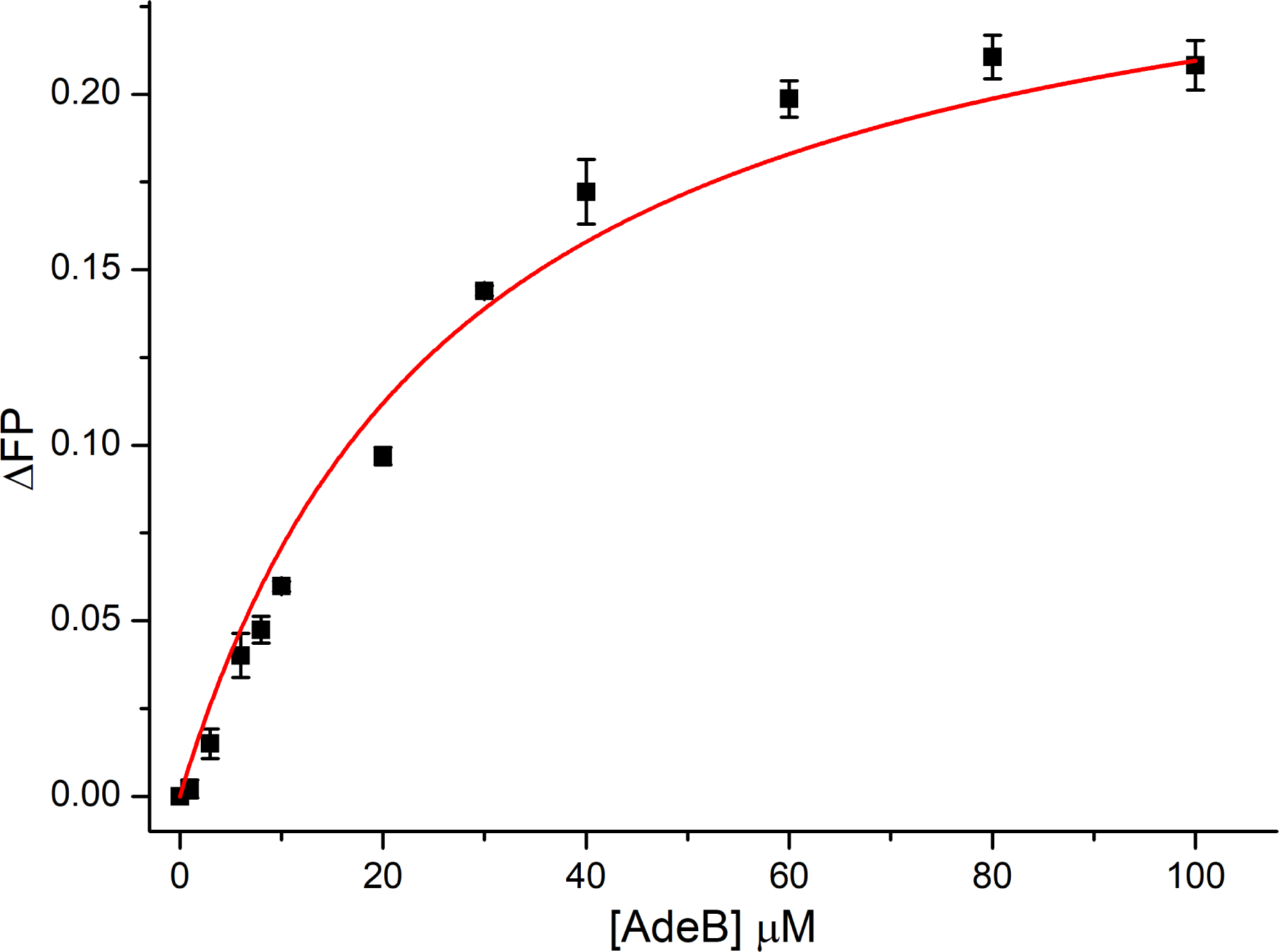
Binding affinity of amotosalen and AdeB determined using fluorescence polarization. Indicated concentrations of AdeB were mixed with 3µM amotosalen. The change in fluorescence polarization signal (ΔFP) indicates a K_D_ of 27.9±1.8 µM for amotosalen.

The previously described cryo-EM structure of AdeB identified a pathway for efflux pump substrate extrusion with entrance through a cleft in the periplasmic domain and sequential binding to proximal and distal multidrug-binding sites (47, 48). Computer modeling of the molecular docking of amotosalen demonstrated binding to both the proximal and distal multidrug bindings sites within the AdeB periplasm domain (see Fig. 2). AMT and 8-MOP were also found to dock with the proximal and distal binding sites, and cleft and distal binding sites, respectively (data not shown). These data are consistent with binding of known efflux substrates to AdeB and homologous RND pumps (31).

**Figure 2.**
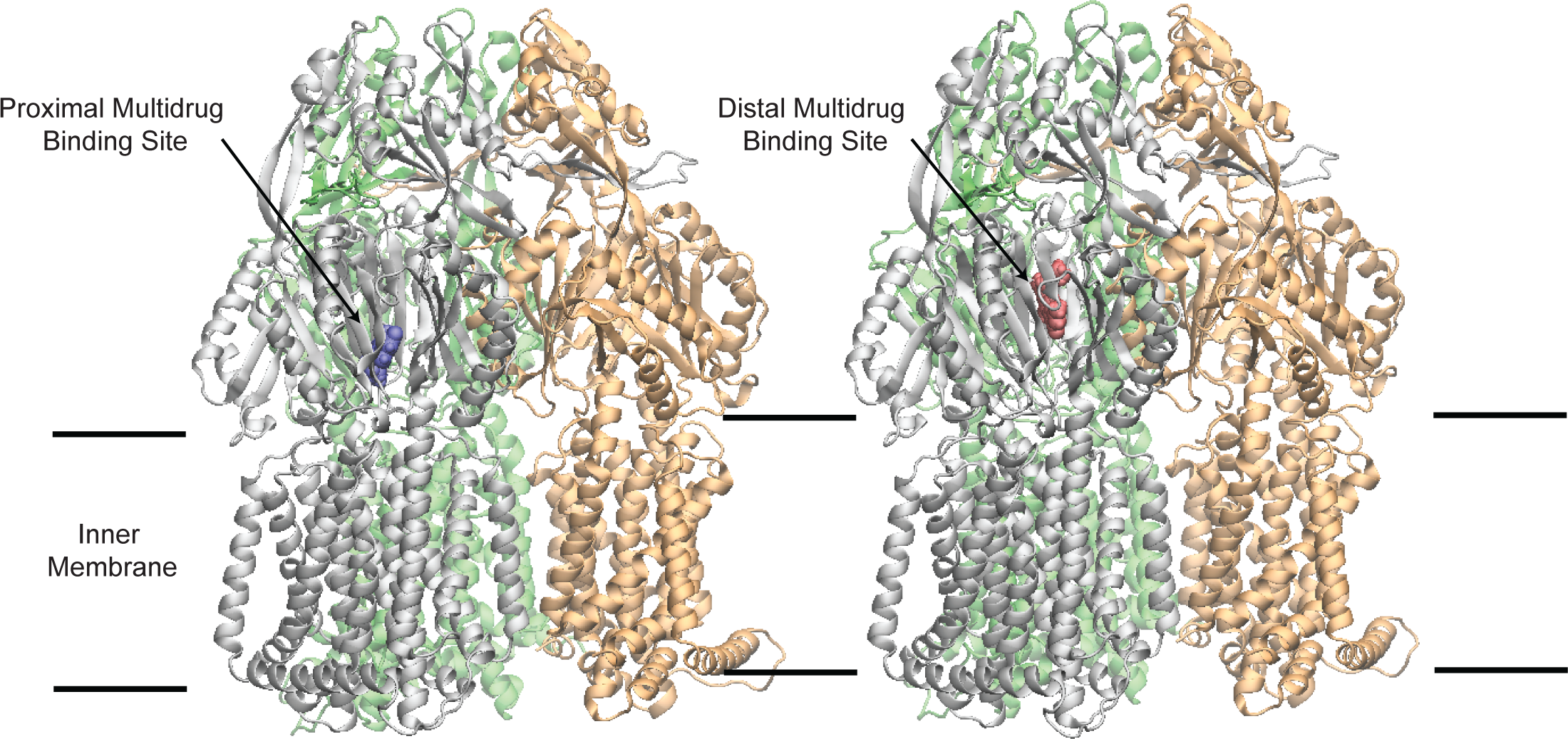
Modeling of amotosalen binding to AdeB. Two amotosalen docking sites were identified in the periplasmic domain of the recently determined cryo-EM structure of AdeB (48) using Induced Fit Docking and XP scores for ranking. The sites are within the hydrophobic cleft between PC1 and PC2 subdomains in AdeB that are thought to form the entry site and pathway for efflux from the periplasm (47). Amotosalen is predicted to bind proximal and distal drug binding sites, as previously defined through autodocking and experimental analysis of antibiotic efflux substrates for AdeB and homologous RND transporters.(47, 83, 84) Specifically, critical contacts are made with hydrophilic residue D664 in the PAIDELGT sequence defined “F-Loop” which forms both part of the cleft entrance and the proximal drug binding site. Amotosalen also binds to the distal multidrug binding site inclusive of hydrophobic interactions with F178, I607 and W610. This hydrophobic patch in the homologous AcrB is critical for stable binding of all substrates, and is further highlighted in corresponding residue interactions in the AcrB-minocycline crystal structure and corresponding MtrD-erythromycin cryo-EM structure (47, 83–85). The predicted binding affinities (XP scores) to the two sites are −10.63 and −11.66, respectively.

## Discussion

Microbial contamination of blood products remains a critical transfusion safety issue. Numerous studies have established the use of amotosalen combined with UVA treatment for broad-spectrum inactivation of bacterial, protozoan, and viral pathogens (27-29, 49-52). Recently, a case of a septic transfusion reaction was reported with a pathogen-reduced product. While a follow-up study indicated that the implicated strains were inactivated by amotosalen (30), this event paired with potential gaps in the existent literature led us to investigate the possibility that MDR Gram-negative organisms, such as *Acinetobacter baumannii*, may have the potential to escape pathogen inactivation.

Gram-negative pathogens in particular are known to have a significant penetration barrier to antimicrobials based on their double cell membrane along with a plethora of multidrug efflux pumps that limit access to the bacterial cytoplasm (53). Here we demonstrate that psoralens including amotosalen are multidrug efflux substrates. It is of interest that the derivation of amotosalen and AMT included the instillation of a primary amine into an existing planar structure with low globularity and few rotatable bonds (Fig. 3). These features taken together are now known to be associated with enhanced penetration of antibiotics across the Gram-negative membrane barrier (54). Therefore, the increased activity (lower MIC values) of amotosalen and AMT compared with 8-MOP is fully consistent with our current understanding of antimicrobial penetrance into Gram-negative pathogens. Nevertheless, access to an intracellular target (in this case, DNA) is a balance of penetration and efflux, and, based on our data, psoralens including amotosalen, appear especially vulnerable to efflux.

**Figure 3.**
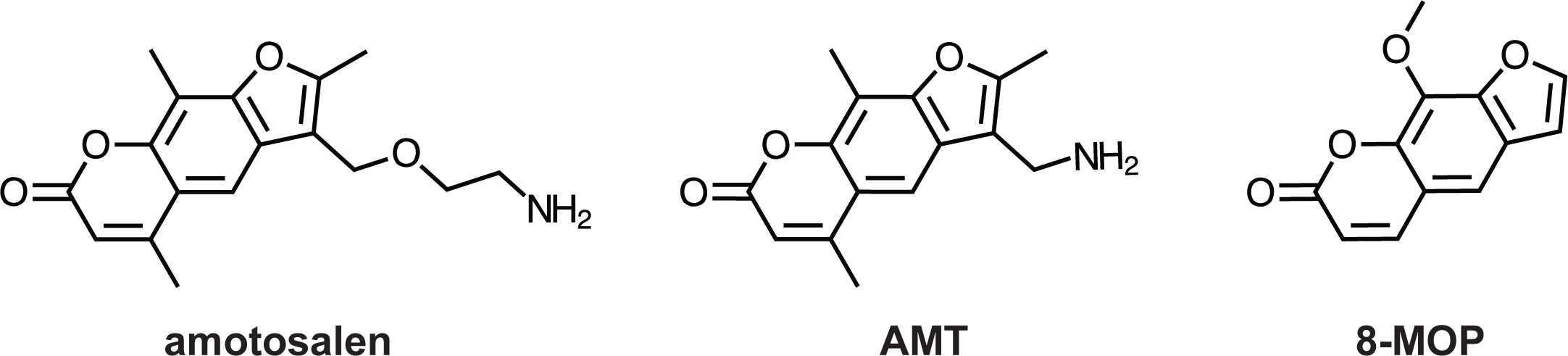
Chemical structures of amotosalen, AMT, and 8-MOP. These related psoralens share a core planar structure with few rotatable bounds. However, amotosalen and 4’-aminomethyltrioxsalen (AMT) also contain primary amino groups, which when combined with molecular planarity and small numbers of rotatable bonds, are associated with enhanced penetrance into Gram-negative pathogens (54). Therefore, structural properties appear to explain the observed enhanced activity of amotosalen and AMT compared with 8-MOP.

Specifically, amotosalen, AMT, and 8-MOP were found to be substrates for the major efflux pumps in *Enterobacterales* and *A. baumannii*. These RND efflux systems, AcrAB-TolC and AdeABC, respectively, share similar substrate specificity and 50% amino acid identity (55). Therefore, the finding that both pumps efflux psoralens, manifest as large increases in the MIC in pump-competent compared with pump-defective strains, is not surprising. Binding of amotosalen to AdeB showed a micromolar dissociation constant, consistent with known affinities of antibiotics to RND pumps (31) and MATE family transporters (56). This affinity is likely located within an optimal dissociation continuum that allows for the ability to bind to several sites within the pump itself with eventual handoff to the outer membrane pump protein and release into the extracellular space (47). Molecular docking to the previously determined cryo-EM structure of AdeB suggest that amotosalen binding is consistent with what has been either determined or predicted for other canonical efflux substrates.

Notably, both AcrABC and AdeABC are upregulated in resistant pathogens. For example, *acrABC* is upregulated in *Enterobacter* species resistant to colistin (57); *E. coli r*esistant to multiple antibiotics (58–60); and in drug-resistant *Salmonella* (61, 62) and *Klebsiella* (63, 64). The expression of *adeABC* is often upregulated in resistant *Acinetobacter* strains (42–44). Therefore, we would expect resistance to amotosalen to be correspondingly increased in these strains.

We found in activity spectrum studies that multidrug-resistant *E. coli, K. pneumoniae, and A. baumannii* demonstrated varying levels of resistance to amotosalen with a significant fraction of isolates with MICs exceeding the concentration of amotosalen used in the Intercept pathogen inactivation procedure. Although we cannot, without extensive genetic and expression analysis, definitively conclude that higher MIC values in these strains were due to efflux activity alone, it is reasonable to hypothesize that efflux is a major contributor. Unlike the pan-susceptible ATCC strains of *E. coli* and *K. pneumoniae* examined, which had low MICs, the otherwise broadly susceptible *A. baumannii* 19798 is known to express AdeABC and the related AdeFGH and AdeIJK pumps potentially explaining its higher intrinsic resistance (65).

We expected that *Pseudomonas aeruginosa*, *Stenotrophomonas maltophilia*, and *Burkholderia* spp. are to be likely candidates for efflux mediated resistance to psoralens, as these bacteria often express or overexpress multiple efflux pumps, resulting in characteristic intrinsic resistance to many antibiotics (66–68). In examination of a limited number of strains (five, two, and eight for these species groups, respectively), we found that they had very high MICs values, often exceeding 256 µM, the highest concentration of amotosalen tested (see Table S1), as well as the 150 µM concentration used during pathogen inactivation of blood products. In the context of the current study, however, we did not examine whether this intrinsic resistance was specifically associated with efflux pump activity, a goal of future work. Interestingly, a minority of strains were either always or variably killed by UVA light in the absence of amotosalen exposure (Table S1). Based on prior literature, we speculate this may result from free radical generation during UVA excitation of bacterial fluorescent pigments and/or endogenous photosensitizers (69, 70).

Though generally effective against most pathogens, psoralens are ineffective against non-enveloped viruses such as HAV, HEV, parvovirus B19, and poliovirus, and relatively imporous bacterial spores (71–73). Taken together, our data now raise the possibility that contemporary multidrug-resistant bacterial isolates have reduced susceptibility to inactivation based on their ability to efflux psoralens and thereby avoid UVA-catalyzed nucleic acid damage. Collectively, our data also suggest the need for further study of psoralen-efflux pump interaction and that future chemical optimization of pathogen inactivating compounds should specifically explore Gram-negative penetrance in the presence of efflux pumps.

Our study has several limitations and results should not be extrapolated directly to the performance of the Cerus INTERCEPT system. Notably, we did not employ the INTERCEPT Illuminator for UVA exposure. We also tested inactivation of pathogens in standard antimicrobial susceptibility testing medium and in lid-less microwell plates with a very short path length for UVA exposure, not in blood products contained in bags with potentially greater UVA opacity. Therefore, our results may differ from the INTERCEPT system when used according to manufacturer’s specifications. Also, our results should be taken in context. MDR Gram-negative pathogens thus far are relatively rare causes of transfusion associated sepsis. In addition, pathogen inactivation strategies have provided significant benefit in reducing the overall frequency of transfusion-associated blood stream infection (74). Nevertheless, emerging antimicrobial resistance, including resistance associated with efflux mechanisms, is becoming increasingly common. Our findings serve as an alert to a potential vulnerability in pathogen inactivation methods that may explain some instances of pathogen inactivation breakthrough and should be an area of further research.

## Materials & Methods

### Chemicals

Amotosalen HCl 3mM solution was obtained from the Cerus INTERCEPT Blood System for Platelets Pathogen Reduction System Dual Storage Processing Set and stored in light-protected aliquots at 4°C. 4’-aminomethyltrioxsalen hydrochloride (AMT) was from Cayman Chemical (Ann Arbor, MI); 8-methoxypsoralen (8-MOP) was from Sigma-Aldrich (St Louis, MO). 8-MOP and AMT were dissolved in DMSO and stored as aliquots at −80°C prior to use.

### Bacterial Strains

Clinical bacterial strains are listed in Table S1 and were obtained from the American Type Culture Collection (ATCC) (Manassas, VA), the CDC-FDA Antimicrobial Resistance Isolate Bank (ARIB) (Atlanta, GA), Walter Reed Army Institute of Research (WRAIR) (Silver Spring, MD) and BEI Resources (Manassas, VA). The *Keio* strain BW25113, and isogenic, JW5503-KanS Δ*tolC E. coli* were obtained from the Coli Genetics Stock Center (Yale University, New Haven, CT).(75) *E. coli* AG100AX Δ*acrAB* Δ*acrEF*(76) was from Ed Yu (Case Western University, Cleveland, OH).

### Creation of isogenic AdeABC efflux strains

Vectors for regulated expression of *adeAB* and *adeC* were created as follows: To create the isopropyl β-D-1-thiogalactopyranoside (IPTG)-inducible, pAdeC vector, pBMTL-2(77) was first converted to pBMTL-2NTC by replacing the kanamycin resistance gene was with a nourseothricin acetyltransferase resistance gene. Specifically, pBMTL-2 was amplified by PCR using “F pLAC (NAT)” and “R pLAC (Nat)” primers (see Table S2), and the nourseothricin resistance gene was amplified from plasmid pMOD3-mNeptune2-nat (78) (Addgene plasmid #120335; http://n2t.net/addgene:120335; RRID:Addgene 120335) using primers “F Nat” and “R Nat” with inclusion of 5’ tails encoding overlap between vector and nourseothricin amplicons, respectively. pBMTL-2 was a gift from Ryan Gill (Addgene plasmid # 22812; http://n2t.net/addgene:22812; RRID:Addgene_22812). All amplification reactions were performed using Q5 high-fidelity DNA polymerase (New England Biolabs, Beverly, MA). PCR amplification was followed by DpnI digestion for 60-90 minutes at 37°C. PCR products were column-purified (Qiaquick PCR Purification kit, Qiagen, Valencia, CA) and assembled using the HiFi reaction kit (New England Biolabs). Transformants were selected on 50 µg/ml nourseothricin. The *adeC* gene from *A. baumannii* strain AYE (ATCC BAA-1710) was then similarly amplified from genomic DNA prepared with the Wizard Genomic DNA Extraction Kit (Promega, Madison, WI) using primers “F AdeC” and “R AdeC” and cloned downstream from the vector pLac site and Shine-Delgarno sequence in pBMTL-2NTC using vector primers, “F pLAC (AdeC)” and “R pLAC (AdeC)”, with the new vector again assembled using HiFi as described above.

To create the arabinose-inducible pAdeAB vector, pBAD-LSSmOrange(79) was amplified using primers “R pBAD” and “F pBAD” to exclude the existing fluorescent protein. pBAD-LSSmOrange was a gift from Vladislav Verkhusha (Addgene plasmid # 37129; http://n2t.net/addgene:37129; RRID:Addgene_37129). *adeAB* genes were amplified from *A. baumannii* AYE genomic DNA, while adding a Shine-Delgarno sequence with optimized spacing from the start codon using primers “F AdeA” and “R AdeB”. Amplicons were assembled by HiFi.

Plasmid constructs were confirmed by Sanger sequencing. Vectors, pAdeC and/or pAdeAB, were introduced into chemically-competent *E. coli* strain AG100AX using the 1X TSS method (80).

### MIC determination

For MIC determination of ATCC, FDA-CDC ARIB, WRAIR, BEI Resources, and *ΔtolC* isogenic strains, bacterial stocks frozen at −80°C were streaked onto tryptic soy agar plates containing 5% sheep blood (Remel, Lenexa, KS) and grown overnight at 35°C in ambient air. Colonies were then suspended in 0.9% saline to 0.5 McFarland, measured using a Densicheck (Biomerieux, Durham, NC); diluted 1:300 in cation-adjusted Mueller-Hinton broth (BD, Franklin Lakes, NJ); and dispensed with an Integra ASSIST (Integra LifeSciences, Plainsboro Township, NJ) into 384-well polystyrene plates (Greiner Bio-One, Monroe, NC) at 50 µL per well.

For MIC determination of AG100AX strains containing pAdeC and pAdeAB plasmids, –80°C frozen stocks were inoculated directly into non-cation-adjusted Mueller-Hinton broth containing 100 µg/ml ampicillin, 50 µg/ml nourseothricin, 5mM calcium chloride, and 5 mM magnesium chloride, with or without *adeABC* induction using 1% L-arabinose and 1.0 mM IPTG (Isopropyl β-D-1-thiogalactopyranoside), and grown overnight at 35°C in 15mL conical tubes with continual rotation. Bacterial cultures were then adjusted to 0.5 McFarland and diluted 1:300 in the same medium, and dispensed as above into microplates.

Filled 384-well plates were centrifuged at 1250 RCF for four minutes to ensure the entire inoculum was in continuity with the bottom of the well. Then two-fold doubling dilutions of stock solutions of amotosalen supplemented to 0.3% Tween-20 (Sigma-Aldrich), AMT, or 8-MOP were dispensed into microwells using the HP D300 digital dispensing system (HP, Inc. Palo Alto, CA), as previously described,(34–37) and the microplates were mixed for 5 minutes on a microplate shaker to ensure complete mixing of psoralens with the inoculum. Microplates were then exposed to ultraviolet light in a UV Stratalinker 1800 (Stratagene, La Jolla, CA), retrofitted with UV BL F8T5 CFL 12-inch UVA 365nm Blacklight Bulbs (Coolspider, Jiinyun, China) and calibrated according to manufacturer’s instructions with an UVA365 UV Light Meter (Amtast, Lakeland, FL).

After incubation at 35°C for 20 h, the A_600_ of the microplate was measured using a TECAN M1000 microplate reader. Minimal inhibitory concentration was determined based on a growth inhibition A_600_ cutoff of 0.06, consistent with our prior determination of absorbance cutoffs for accurate MIC determinations in 384-well plate format (35, 37, 81). Notably, the AdeABC pump was inactive in *E. coli* AG100X in Mueller-Hinton broth without the addition of divalent cations, consistent with prior use of magnesium supplementation when expressing this *A. baumannii* efflux pump in *E. coli* (82). The concentrations of MgCl_2_ and CaCl_2_ used were empirically determined to be optimal for efflux of minocycline and ethidium bromide (data not shown).

### AdeB binding affinity and molecular docking

His-tagged AdeB protein was purified as described previously (31). Fluorescence polarization assays were performed in a ligand binding solution consisting of 20 mM HEPES-NaOH pH7.5 and 0.05% n-dodecyl-β-D-maltoside (DDM) and 3 μM amotosalen. The experiments were done by titrating the AdeB protein in solution containing 20 mM HEPES-NaOH pH7.5 and 0.05% DDM into the ligand binding solution while keeping DDM concentration constant. Fluorescent polarization was measured at 25°C using a PerkinElmer LS55 spectrofluorometer coupled with a Hamamatsu R928 photomultiplier. The excitation wavelength for amotosalen was 350 nm and the fluorescent polarization signal (ΔP) was measured at 470 nm. Titration data points represent 15 measurements and 3 biological replicates were performed to determine the K_D_ as previously described (31). ORIGIN Version 7.5 (OriginLab Corp., Northampton, MA) was used for curve fitting.

### Molecular Docking

A structure of the “binding” protomer of AdeB-Et-I (PDB ID: 7KGH) was used to as the template, in which the bound Et ligands were removed from the protomer (48). The Protein Preparation Wizard Module of Maestro (Release 2019-3) (Schrödinger, New York, NY) was used for induced-fit-docking simulations^46^ using default parameters to predict binding modes of amotosalen, AMT, and 8-MOP to AdeB. For each calculation, residues within 5Å of the bound ligand were selected for side chain optimization using Prime refinement. The docking results with the lowest XP scores were selected as predicted poses.

## Acknowledgements

This work was supported by R01AI145069 to E. Yu and R21AI146485 to J.E.K. K.E.Z. was supported in part by a National Institute of Allergy and Infectious Diseases training grant (T32AI007061). W.A.F. was supported by a NIH Clinical Center Intramural Research Program grant (ZIA CL002128). The content is solely the responsibility of the authors and does not necessarily represent the view of the National Institutes of Health, the Department of Health and Human Services, or the U.S. Federal Government. The HP D300 digital dispenser and TECAN M1000 used in experiments were provided by TECAN (Morrisville, NC). TECAN had no role in study design, data collection/interpretation, manuscript preparation, or decision to publish.

